# Plant silicon defence disrupts cryptic colouration in an insect herbivore by restricting carotenoid sequestration into the haemolymph

**DOI:** 10.1101/2025.04.02.646851

**Authors:** Tarikul Islam, Sidra Anwar, Christopher I. Cazzonelli, Ben D. Moore, Scott N. Johnson

## Abstract

1. Cryptic colouration is a primary anti-predation strategy in herbivorous insects. Achieving crypsis often requires acquiring dietary carotenoids—tetraterpene pigments vital for plant colouration and photoprotection. Silicon (Si) accumulation in plants makes tissues tougher and less digestible for insects, but its effect on plant pigment levels remains unclear. It is also unknown whether feeding on silicified plants impairs insects’ ability to sequester carotenoids and achieve crypsis.
2. Using the model grass *Brachypodium distachyon*, we demonstrate that the cotton bollworm (*Helicoverpa armigera*) larvae exhibited stunted growth and reduced carotenoid sequestration, particularly lutein, into their hemolymph when feeding on Si-supplemented (+Si) plants. This reduction led to distinct body-colour morphs: larvae feeding on +Si plants developed brown colouration, contrasting sharply with the green leaves, whereas larvae feeding on Si-free (−Si) plants exhibited green cryptic colouration that blended seamlessly with the foliage.
3. Plant leaves contained various carotenoids (neoxanthin, violaxanthin, β-carotene and lutein) and chlorophylls (a and b), but larvae only sequestered β-carotene and lutein into their haemolymph while excreting substantial amounts of pigments, regardless of plant Si status. Under insect-free conditions, +Si plants had lower carotenoid and chlorophyll contents than −Si plants. However, following insect herbivory, pigment levels in −Si and +Si plants equalised. Insect herbivory also increased Si accumulation in +Si plants.
4. Our findings provide novel evidence that plant Si defences can disrupt cryptic colouration in insect herbivores by restricting carotenoid sequestration from host plant tissues. This disruption could increase insect visibility to predators, potentially elevating their risk of predation.

## INTRODUCTION

Natural enemies of insect herbivores, such as predators and parasitoids, locate their prey by exploiting optical (e.g. prey colour, shape, size) and chemical cues (e.g. HIPVs, prey odours, pheromones) (Dicke, 2009; Harmon et al., 1998; Vosteen et al., 2016). Many predators, including vertebrates (e.g. birds, amphibians) and invertebrates (e.g. spiders, ground beetles, ants), rely primarily on vision to hunt insect prey (Gonzalez-Bellido et al., 2022; Lim & Ben-Yakir, 2020). As a counteradaptation, some insects have evolved cryptic colouration, a camouflage strategy where they adopt the colours of their surroundings to blend into the environment (Greeney et al., 2012). Crypsis can reduce the probability of being detected and killed by predators reliant on vision, particularly when insects exhibit crypsis against a uniform background (Lyytinen, 2001).

Carotenoids, a class of lipophilic tetraterpene pigments (C_40_), are often essential for the cryptic colouration of insect bodies and eggs (Heath et al., 2012). Except for some aphids, insect herbivores cannot synthesise their own carotenoids and typically obtain them from their diet (Maoka, 2020; Moran & Jarvik, 2010). For example, lepidopteran larvae sequester carotenoids, such as lutein and β-carotene, derived from their host plants into their haemolymph to produce green cryptic colouration (Grayson et al., 1991; Hackman, 1952). This protective colouration allows larvae, particularly in later instars, to blend with green leaves and evade predation (Czeczuga, 1986; Nguyen et al., 2019). The green haemolymph, often termed insectoverdin, is a mixture of yellow and blue chromoproteins. The yellow chromoprotein contains carotenoids (i.e. lutein or β-carotene, or both) as prosthetic groups and the blue chromoprotein includes bile pigments, such as pterobilins and mesobiliverdin (Hackman, 1952; Heath et al., 2012). Larvae can derive these blue pigments by breaking down plant chlorophylls or synthesising them independently (Hackman, 1952; Welch et al., 2017). Hence, larval green colouration largely depends on acquiring carotenoids from host plant tissues. Failure to sequester lutein or β-carotene can disrupt larval crypsis, increasing their risk of predation (Nguyen et al., 2019; Valadon et al., 1974; Welch et al., 2017).

The acquisition of carotenoids by larvae depends on the carotenoid contents of their host plants (Eichenseer et al., 2002; Heath et al., 2012). Plants produce carotenoids as pigments that colour tissues (yellow to red), harvest light energy for photosynthesis alongside chlorophylls (green pigments) and protect against photooxidative damage, including that caused by insect herbivory (Baranski & Cazzonelli, 2016; Pérez-Gálvez et al., 2020; Schaefer & Rolshausen, 2006). Plant leaves contain several carotenoids, of which violaxanthin, β-carotene, lutein and neoxanthin are usually the most abundant (Cazzonelli, 2011; Heath et al., 2012). Insect herbivory can alter pigment biosynthesis in plants (Bi & Felton, 1995; Schaefer & Rolshausen, 2006). In particular, oxidative stress from insect feeding can lead to cleavage of carotenoids into volatile chemicals that deter insects (Meng et al., 2023).

Grasses, such as cereals, are hyper-accumulators of silicon (Si), which can account for up to 10% of their shoot dry weight and are reliant on Si defences to combat herbivore attack (Hodson et al., 2005; Vicari & Bazely, 1993). Si uptake from the soil and deposition in plant tissues as silica (SiO_2_) confers physical resistance to insect herbivores (Epstein, 2009). Increasingly, studies show that Si impacts a broader range of anti-herbivore defences in plants, including secondary metabolite defences that act directly (Epstein, 2009; Reynolds et al., 2009) or indirectly via herbivore-induced plant volatiles that recruit natural enemies (Islam et al., 2022b). Given that lepidopteran larvae feed less on silicified plants and silicification reduces the digestibility of ingested tissues (Hunt et al., 2008; Massey & Hartley, 2009), we reason that larvae may accumulate less carotenoids when feeding on Si-supplemented plants, potentially disrupting their green colouration. Furthermore, both Si accumulation and pigment production can be induced as part of plant defence responses (Islam et al., 2020; Schaefer & Rolshausen, 2006; Waterman et al., 2020); yet it remains unknown whether Si supplementation impacts the levels of carotenoids and chlorophylls in plant tissues under insect herbivory, nor how this affects carotenoid acquisition in insect larvae.

Using *Brachypodium distachyon*, a model grass (Girin et al., 2014), we investigated the plant-mediated effects of Si on pigment sequestration and cryptic colouration of a generalist chewing insect, the cotton bollworm, *Helicoverpa armigera* (Lepidoptera: Noctuidae). *Helicoverpa armigera* is a cosmopolitan pest of numerous crop plants, including economically important Triticeae crops (Cunningham & Zalucki, 2014), that have a close phylogenetic connection to *B. distachyon* (Brkljacic et al., 2011). Studies have documented that *H. armigera* larvae actively feed on *B. distachyon* under controlled conditions (Biru et al., 2021; Islam et al., 2022a; Waterman et al., 2020). While larvae prefer to feed on plant reproductive structures (i.e. flowers and fruits), they also feed on leaves (Rogers & Brier, 2010; Zalucki et al., 1986). Larval feeding on different host plants or plant parts often leads to body-colour polymorphism, conspicuously in later instars (Yamasaki et al., 2009). Specifically, larvae are more likely to exhibit green colouration when feeding on green leaves (Yamasaki et al., 2009).

Our objectives were to (i) investigate the plant-mediated effects of Si on pigment sequestration into the haemolymph of *H. armigera* larvae and the exhibition of larval green cryptic colouration; and (ii) elucidate the effects of Si and insect herbivory on pigment levels in *B. distachyon* leaves. We hypothesise that Si supply and insect herbivory influence carotenoid and chlorophyll levels in leaf tissues and that larval feeding on silicified leaves restricts carotenoid sequestration into the haemolymph, disrupting larval green colouration.

## MATERIALS AND METHODS

### Plant growth conditions, Si supplementation, and insect rearing

*Brachypodium distachyon* seeds (accession Bd21-3, INRAE, Paris, France) were germinated in irrigated perlite as per the procedures described previously (Hall et al., 2020). Briefly, seeds were soaked in water, manually dehusked using forceps, and sterilised in bleach. After thoroughly washing with water, seeds were sown in a wet perlite medium and cold stratified at 4°C for seven days. Subsequently, seedlings were grown for two weeks to ensure uniform development. Seedlings were transplanted into non-aerated hydroponic systems, consisting of two nested plastic cups (480 ml) and a fitted foam disc at the top to anchor each seedling, as described in Hall et al. (2020). Cups were filled weekly with 370 ml fresh nutrient solution either with or without Si supplementation. The nutrient solution comprised 1 mM KNO_3_, 1 mM Ca(NO_3_)_2_.4H_2_O, 1 mM KH_2_PO_4_, 0.6 mM MgSO_4_, 0.1 mM NaCl, 15 μM H_3_BO_3_, 0.5 μM MnCl_2_.4H_2_O, 0.7 μM ZnSO_4_.7H_2_O, 0.8 μM Na_2_MoO_4_.2H_2_O, 0.8 μM CuSO_4_.5H_2_O and 0.12 mM NaFe(iii)EDTA.

Plants were provided with Si by adding liquid K_2_SiO_3_ (21% K_2_O and 32% SiO_2_, Agsil32, PQ Australia, SA, Australia) to the nutrient solution at 2 mM concentrations (SiO_2_ equivalent). Previous studies have shown that *B. distachyon* can accumulate around 2-4% Si in leaves at this supplementation rate (Hall et al., 2020; Islam et al., 2022a), aligning with Si concentrations reported in other *Brachypodium* species, Poaceous crops and turfgrasses in natural habitats (Guntzer et al., 2011; Hodson et al., 2005). To balance the added K^+^ ions in the +Si solution, KCl (Sigma-Aldrich, MO, USA) was added to the −Si solution, and both solutions were adjusted to pH 5.5 using HCl. Plants were grown under natural lighting in a glasshouse maintained at 22/18°C (day/night) temperatures and 50% (± 6%) relative humidity. Plants were rotated weekly within the glasshouse to minimise positional bias. *Helicoverpa armigera* larvae were obtained from CSIRO Agriculture and Food, Narrabri, NSW, Australia. Larvae were reared on an artificial diet adapted from Teakle and Jensen (1985), consisting primarily of soya flour, wheat germ, brewer’s yeast, agar and distilled water. The diet was supplemented with trace amounts of preservatives, including nipagen, sorbic acid and ascorbic acid.

### Herbivore treatment: larval growth, colour contrast and sampling for pigment analysis

Plants were assigned for herbivore treatment (i.e. insect or insect-free) after five weeks of growth in hydroponic containers. Forty plants (20 +Si and 20 −Si) each received a single *H. armigera* second instar larva, while 20 plants (10 +Si and 10 −Si) remained insect-free. Larvae were starved for 24 hr and weighed before placing on plants. All plants were caged in transparent acrylic cylinders with mesh apertures, as described previously (Johnson et al., 2020). After ten days, larvae were removed from plants and starved individually in Petri dishes for 24 hr to allow for frass evacuation before being weighed. The relative growth rate (RGR) of larvae was calculated as mass gain (mg) relative to initial mass (mg) per day (Massey & Hartley, 2009).

Frass expelled by larvae during the final 48 hrs was collected from hydroponic containers and corresponding Petri dishes. For each Si treatment, frass from 3-4 larvae was combined to create samples of 25-30 mg (*N* = 5-6 per treatment) and stored at −80°C for pigment analysis.

Twenty larvae (10 fed on −Si plants and 10 fed on +Si plants) were randomly selected and photographed using a stereo microscope (Zeiss AX10) equipped with a camera (Zeiss AxioCam MRc). Each larva was placed individually over a leaf from a matching treatment plant (e.g. larvae fed on −Si plants were photographed on −Si leaves). Photos were processed in ImageJ to quantify the contrast between larvae and their green leaf backgrounds. The RGB (red, green and blue) values of larvae (R_1_, G_1_ and B_1_) and leaf backgrounds (R_2_, G_2_ and B_2_) were measured, and colour differences were calculated using the Euclidean distance formula 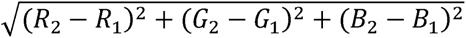 (Ahmad et al., 2019). Higher Euclidean distances indicated greater colour contrast between larvae and their leaf backgrounds.

Larvae were subsequently placed at −20°C for 5 min to restrict movements, and a proleg was removed for haemolymph collection. Haemolymph was also collected from 20 fifth instar larvae reared exclusively on the artificial diet to verify that haemolymph pigments were plant-derived. Haemolymph samples were obtained by pooling 20 µl haemolymph from 2-3 larvae using a sterile micropipette. The samples (*N* = 6-9 per treatment) were flushed immediately into microcentrifuge tubes containing 20 µl of cold phosphate-buffered saline (pH 7.4) and stored at −80°C for further analysis. To assess the effects of Si and insect herbivory on plant pigments, the youngest fully expanded leaves from insect-free and insect-attacked plants (*N* = 4-5 per Si treatment) were harvested. Leaves were snap-frozen in liquid nitrogen and stored at −80°C for subsequent analysis.

### Pigment extraction and quantification

Pigments in leaf and frass samples were extracted following the methods described in Dhami et al. (2018) and Alagoz et al. (2020). For each sample, 25-40 mg leaf tissue or 25-30 mg frass was placed in a microcentrifuge tube and ball-milled using a tissue lysing system (Qiagen, TissueLyser II). Five hundred microliters of extraction buffer (6:4 v/v acetone:ethyl acetate with 0.1% butylated hydroxytoluene) was added to the ground leaf tissue or frass and vortexed for one minute. Subsequently, 500 µl of Milli-Q water was added to the mixture for phase separation and centrifuged at 15000 rpm for 5 min at 4°C. The pigment-rich upper ethyl acetate phase (∼250 µl) was collected into a separate microcentrifuge tube and centrifuged again for 5 min to yield ∼150 µl of purified extract. For haemolymph samples, pigments were extracted similarly, excluding the ball-milling step.

Extracts were stored at 4°C until analysis using an HPLC system (Agilent 1260 Infinity, California, USA) equipped with a diode array detector following Alagoz et al. (2020). Twenty microliters of extract per sample was injected and run through a GraceSmart C18 HPLC column (4.6 mm × 250 mm column, 4 µm size). Pigments were separated using three mobile phase solvents (9:1:0.01 v/v/v acetonitrile:water:triethylamine, 100% ethyl acetate and 100% acetonitrile). Identification of pigments was based on their retention times relative to synthetic standards and absorption spectra at 440 nm. Pigment contents were quantified using standard curves derived from peak areas and expressed as micrograms per gram of sample fresh weight for leaf and frass samples and as micrograms per millilitre of sample volume for haemolymph (Anwar et al., 2022).

### Si analysis in leaf tissues

Concentrations of total leaf Si (% dry weight) were quantified in 20 +Si plants (10 insect-fed and 10 insect-free) and 10 −Si plants (five insect-fed and five insect-free) as per Reidinger et al. (2012). Plants were harvested and oven-dried at 60°C for seven days, and the leaves were ball-milled to a fine powder. Approximately 80 mg of ground leaf tissue per sample was analysed using an X-ray fluorescence spectrometer (Epsilon 3^x^; PANalytical, EA Almelo, The Netherlands). Spectrometer readings were standardised against a citrus leaf sample of known Si concentration (NCS ZC73018 Citrus leaves, China National Institute for Iron and Steel). Leaf Si concentrations in −Si plants were below the detection limit of the spectrometer (< 0.3%) and were therefore excluded from the analysis.

### Statistical analysis

All data were analysed using R (version 4.3.0) (R Core Team, 2019). Haemolymph samples from larvae reared on the artificial diet lacked any detectable pigments and were therefore excluded from the analysis. The effects of Si and insect attack and their interactions on leaf pigment contents were analysed using two-way ANOVAs from the ‘car’ package in R. Assumptions of distributions and homogeneity of variances for ANOVAs were verified with QQ plots and residual versus fits plots, respectively. To further compare carotenoid and chlorophyll contents in −Si and +Si leaves under insect-attacked and insect-free conditions, Welch’s *t*-tests were performed, and *p*-values were adjusted for multiple comparisons using the Benjamini-Hochberg method (FDR). All other response variables were analysed using Welch’s *t*-tests. Hedges’ *g* was calculated to measure the effect sizes. One leaf sample from an insect-attacked +Si plant and one frass sample from larvae on −Si plants were excluded from the analysis as extreme outliers.

## RESULTS

### Larval performance, cryptic colouration, pigment sequestration and excretion

Si supplementation reduced larval RGR by 63% (Fig. 1a; Table 1) and increased larval contrast against green leaves by over threefold compared to larvae on −Si plants (Fig. 1b; Table 1).

**Fig. 1.**
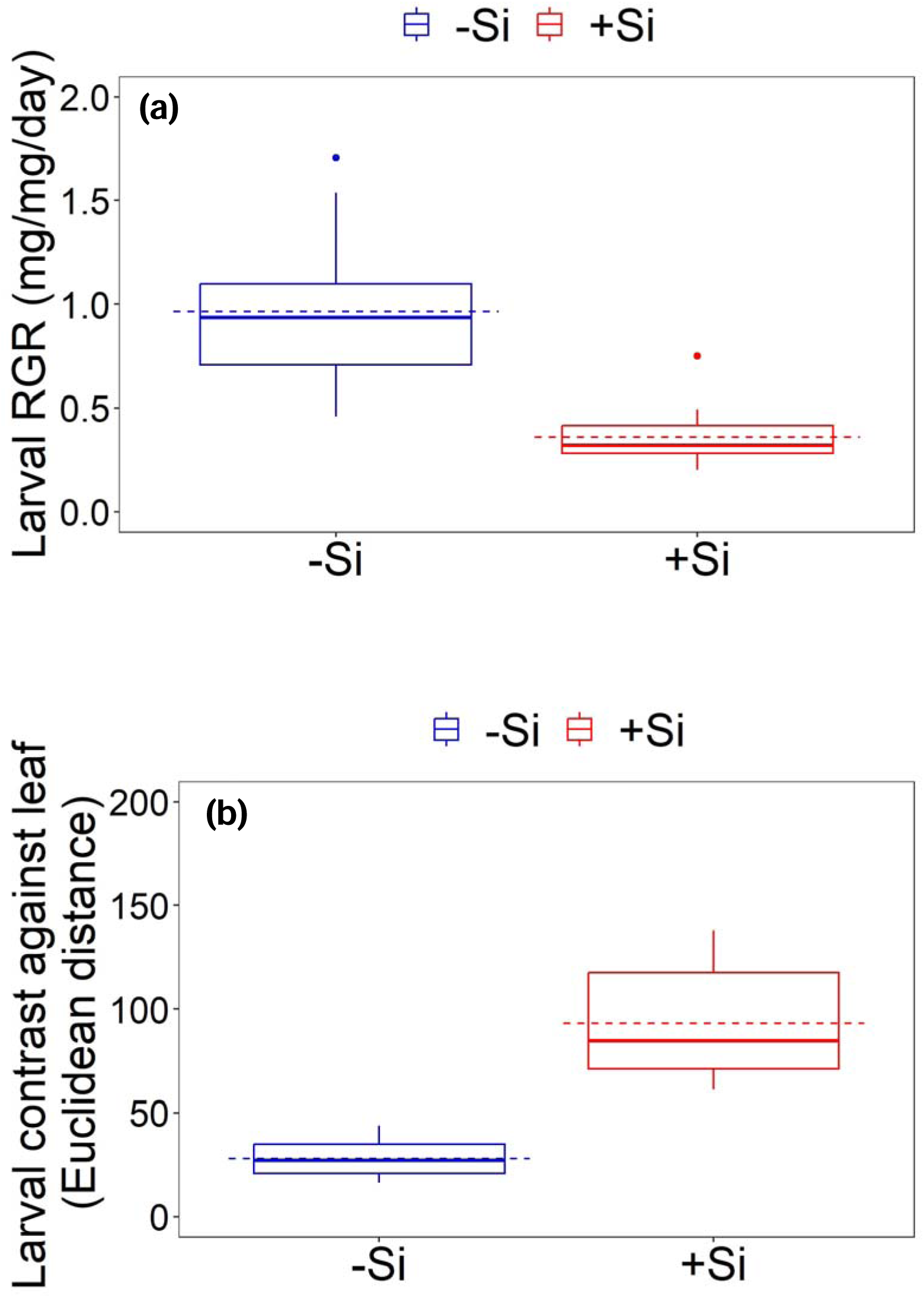
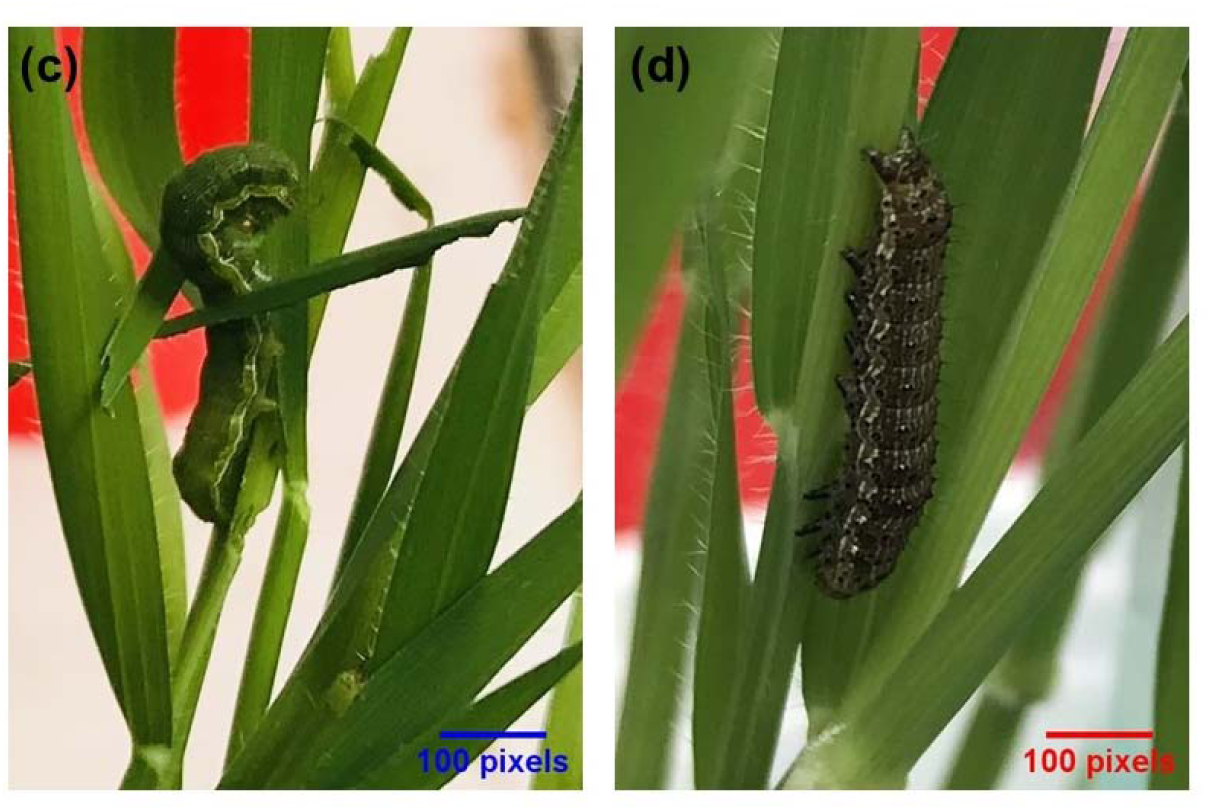
Relative growth rate (mg/mg/day) (a) and colour contrast scores (b) of larvae when feeding on –Si (silicon-free) and +Si (silicon-supplemented) plants. Larvae exhibited distinct colour morphs: cryptic green on −Si plants (c) and conspicuous brown on +Si plants (d). Boxplots show the interquartile ranges (boxes), non-outlier ranges (whiskers), outliers (solid dots), medians (solid lines) and means (dashed lines). Treatment means were compared using Welch’s *t*-tests. Asterisks indicate levels of statistical significance (*** *p* < 0.001) at a 95% confidence level.

**Table 1.**
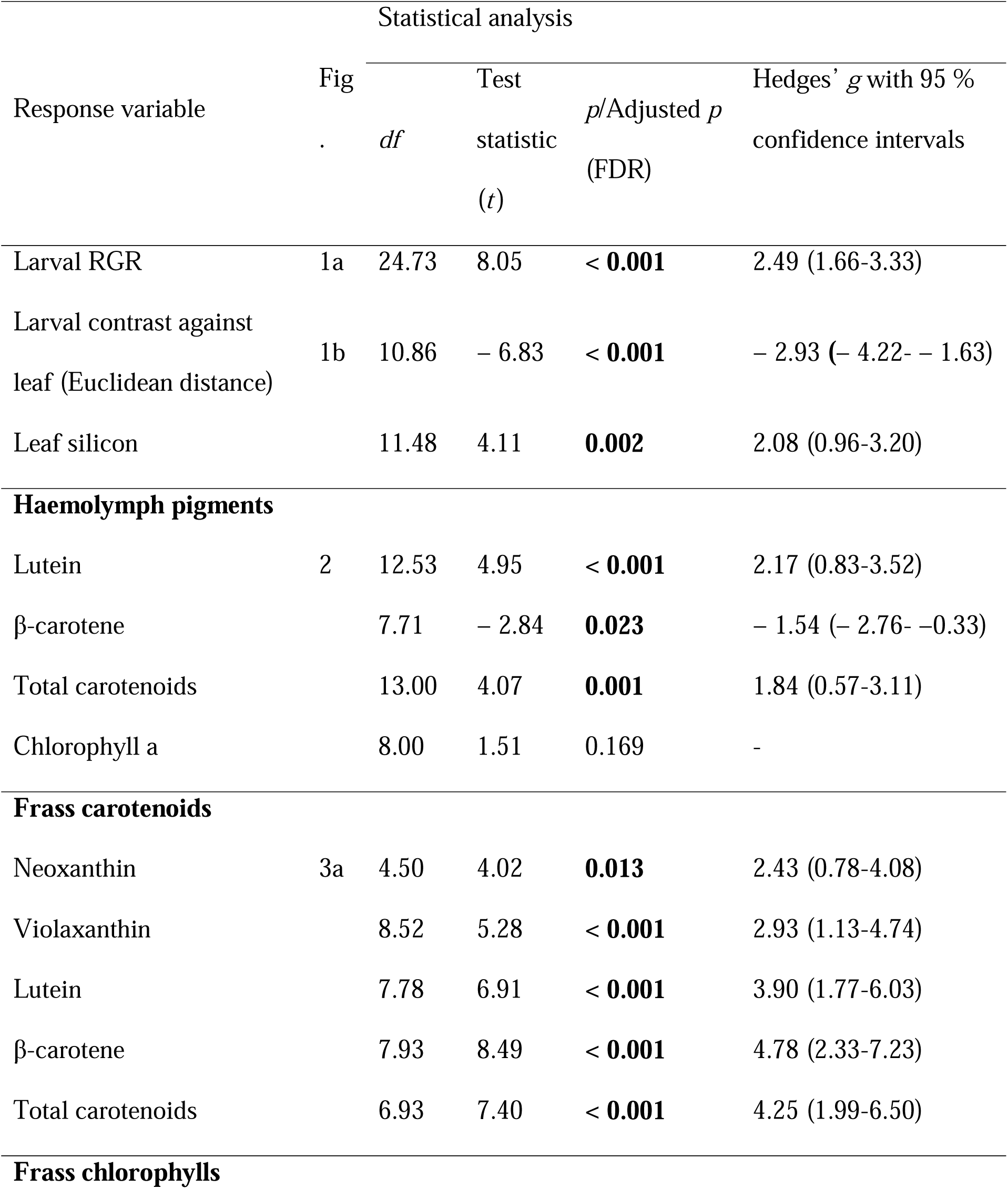
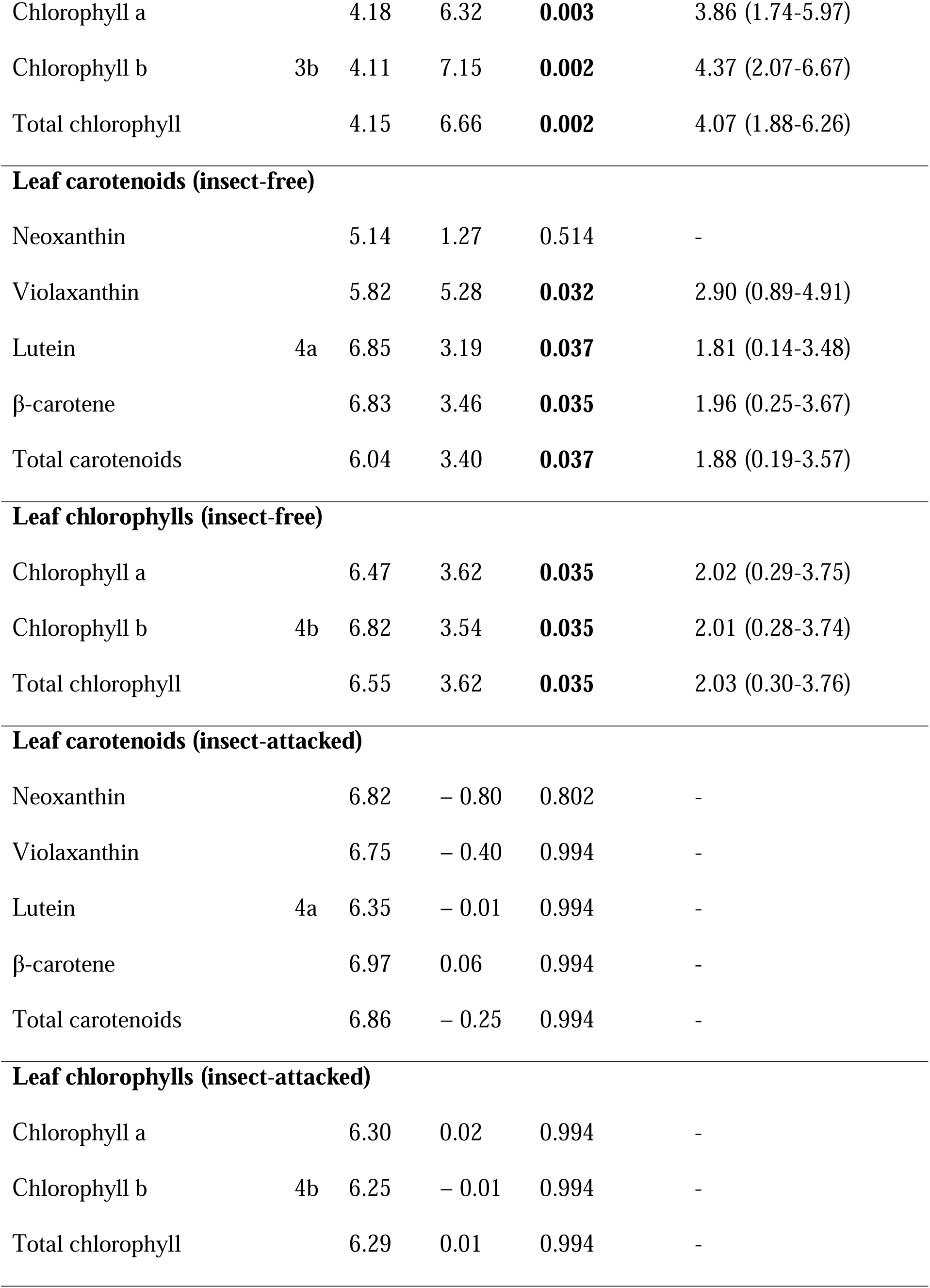
Results of Welch’s *t*-tests and effect size estimates (Hedges’ *g*). Statistically significant effects (*p* < 0.05) are indicated in bold. Effect sizes greater than 0.8 (absolute value) are considered large.

| Response variable | Fig | Statistical analysis |  |  | Hedges' *g* with 95 % confidence intervals |
| --- | --- | --- | --- | --- | --- |
|  |  | *df* | Test statistic (*t*) | *p*/Adjusted *p* (FDR) |  |
| Larval RGR | 1a | 24.73 | 8.05 | **< 0.001** | 2.49 (1.66-3.33) |
| Larval contrast against leaf (Euclidean distance) | 1b | 10.86 | – 6.83 | **< 0.001** | – 2.93 (– 4.22- – 1.63) |
| Leaf silicon |  | 11.48 | 4.11 | **0.002** | 2.08 (0.96-3.20) |
| **Haemolymph pigments** |  |  |  |  |  |
| Lutein | 2 | 12.53 | 4.95 | **< 0.001** | 2.17 (0.83-3.52) |
| β-carotene |  | 7.71 | – 2.84 | **0.023** | – 1.54 (– 2.76- – 0.33) |
| Total carotenoids |  | 13.00 | 4.07 | **0.001** | 1.84 (0.57-3.11) |
| Chlorophyll a |  | 8.00 | 1.51 | 0.169 | - |
| **Frass carotenoids** |  |  |  |  |  |
| Neoxanthin | 3a | 4.50 | 4.02 | **0.013** | 2.43 (0.78-4.08) |
| Violaxanthin |  | 8.52 | 5.28 | **< 0.001** | 2.93 (1.13-4.74) |
| Lutein |  | 7.78 | 6.91 | **< 0.001** | 3.90 (1.77-6.03) |
| β-carotene |  | 7.93 | 8.49 | **< 0.001** | 4.78 (2.33-7.23) |
| Total carotenoids |  | 6.93 | 7.40 | **< 0.001** | 4.25 (1.99-6.50) |
| **Frass chlorophylls** |  |  |  |  |  |
| Chlorophyll a |  | 4.18 | 6.32 | **0.003** | 3.86 (1.74-5.97) |
| Chlorophyll b | 3b | 4.11 | 7.15 | **0.002** | 4.37 (2.07-6.67) |
| Total chlorophyll |  | 4.15 | 6.66 | **0.002** | 4.07 (1.88-6.26) |
| **Leaf carotenoids (insect-free)** |  |  |  |  |  |
| Neoxanthin |  | 5.14 | 1.27 | 0.514 | - |
| Violaxanthin |  | 5.82 | 5.28 | **0.032** | 2.90 (0.89-4.91) |
| Lutein | 4a | 6.85 | 3.19 | **0.037** | 1.81 (0.14-3.48) |
| β-carotene | | 6.83 | 3.46 | **0.035** | 1.96 (0.25-3.67) |
| Total carotenoids |  | 6.04 | 3.40 | **0.037** | 1.88 (0.19-3.57) |
| **Leaf chlorophylls (insect-free)** |  |  |  |  |  |
| Chlorophyll a |  | 6.47 | 3.62 | **0.035** | 2.02 (0.29-3.75) |
| Chlorophyll b | 4b | 6.82 | 3.54 | **0.035** | 2.01 (0.28-3.74) |
| Total chlorophyll |  | 6.55 | 3.62 | **0.035** | 2.03 (0.30-3.76) |
| **Leaf carotenoids (insect-attacked)** |  |  |  |  |  |
| Neoxanthin |  | 6.82 | - 0.80 | 0.802 | - |
| Violaxanthin |  | 6.75 | - 0.40 | 0.994 | - |
| Lutein | 4a | 6.35 | - 0.01 | 0.994 | - |
| β-carotene | | 6.97 | 0.06 | 0.994 | - |
| Total carotenoids |  | 6.86 | - 0.25 | 0.994 | - |
| **Leaf chlorophylls (insect-attacked)** |  |  |  |  |  |
| Chlorophyll a |  | 6.30 | 0.02 | 0.994 | - |
| Chlorophyll b | 4b | 6.25 | - 0.01 | 0.994 | - |
| Total chlorophyll |  | 6.29 | 0.01 | 0.994 | - |

Larvae feeding on −Si plants exhibited green cryptic colouration, blending with the green leaves, whereas larvae feeding on +Si plants exhibited brown colouration (Fig. 1b). On +Si plants, larvae sequestered lower levels of total carotenoids (−44%) in their hemolymph, which was driven mostly by reduced sequestration of lutein (−53%) (Fig. 2). Sequestration of β-carotene in the haemolymph rose by 58% in larvae feeding on +Si plants compared to those feeding on −Si plants, but β-carotene represented a small proportion of total carotenoids (14% when averaged across Si treatments) (Fig. 2; Table 1). Neoxanthin and violaxanthin were completely absent in the haemolymph, irrespective of larval feeding on −Si or +Si plants. Chlorophyll pigments were generally absent in the haemolymph, except for a few larvae (20%) that sequestered a negligible content of chlorophyll a when feeding on −Si plants (Fig. 2; Table 1). Larvae on the artificial diet did not sequester any carotenoid or chlorophyll pigments and developed brown morphs (Fig. 2), suggesting that haemolymph carotenoids originated from leaves. All larvae feeding on plants excreted carotenoid and chlorophyll pigments in their frass, irrespective of the Si status of their host plants. Larvae feeding on +Si plants excreted 50% less carotenoids in their frass compared to larvae on −Si plants, with levels of neoxanthin (−85%), β-carotene (−59%) and lutein (−45%) being particularly low (Fig. 3a; Table 1). Similarly, the total chlorophyll content of frass was 77% lower when larvae fed on +Si plants, with both chlorophyll a (−78%) and chlorophyll b (−77%) showing a decline (Fig. 3b; Table 1).

**Fig. 2.**
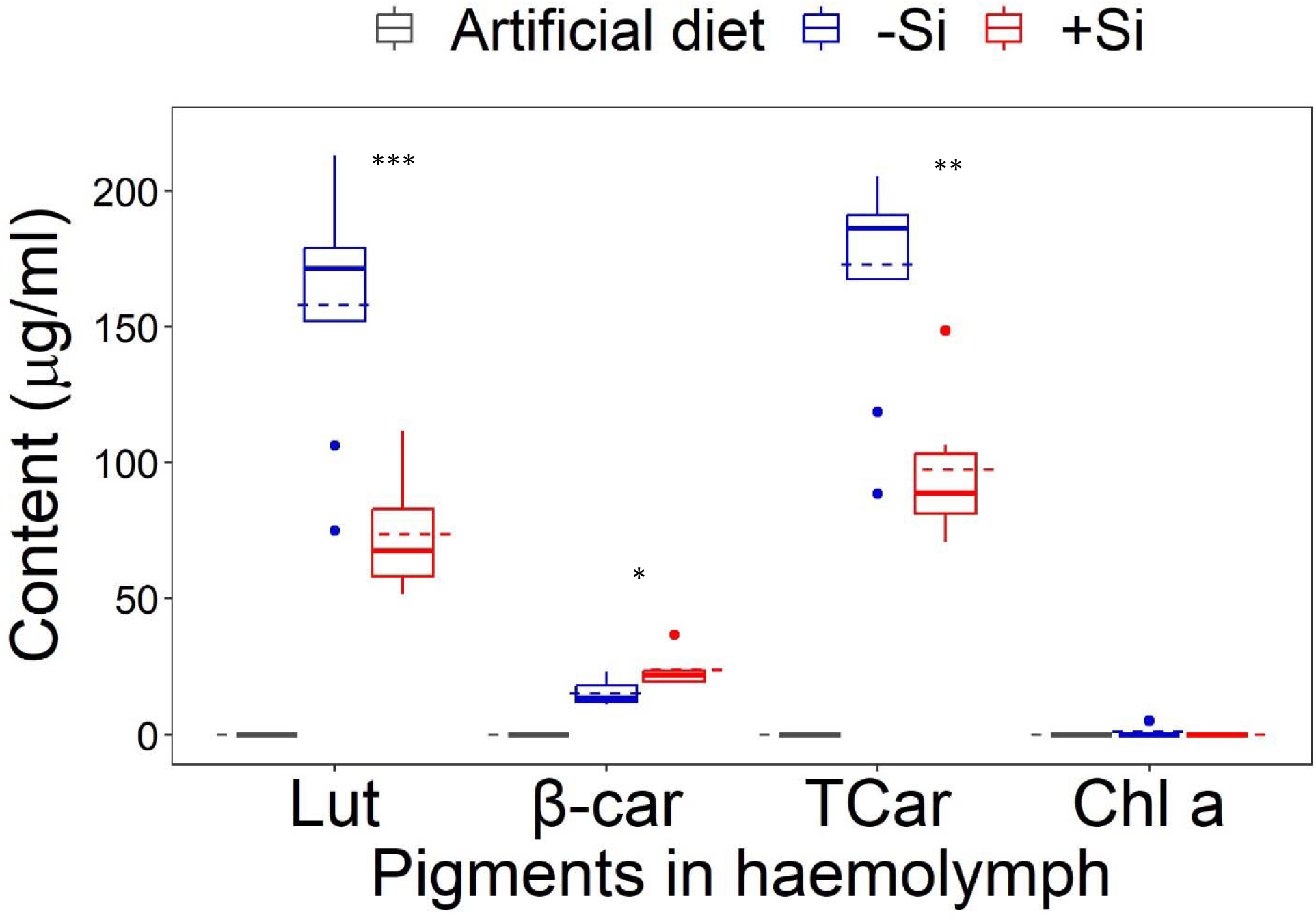
Pigment contents in the haemolymph (µg/ml) of larvae fed on –Si (silicon-free) plants, +Si (silicon-supplemented) plants and an artificial diet. The boxplot shows the interquartile ranges (boxes), non-outlier ranges (whiskers), outliers (solid dots), medians (solid lines) and means (dashed lines). Treatment means (–Si and +Si) were compared using Welch’s *t*-tests. Asterisks indicate levels of statistical significance (* *p* < 0.05, ** *p* < 0.01, *** *p* < 0.001) at a 95% confidence level. Lutein (Lut), β-carotene (β-car), total carotenoids (TCar) and chlorophyll a (Chl a).

**Fig. 3.**
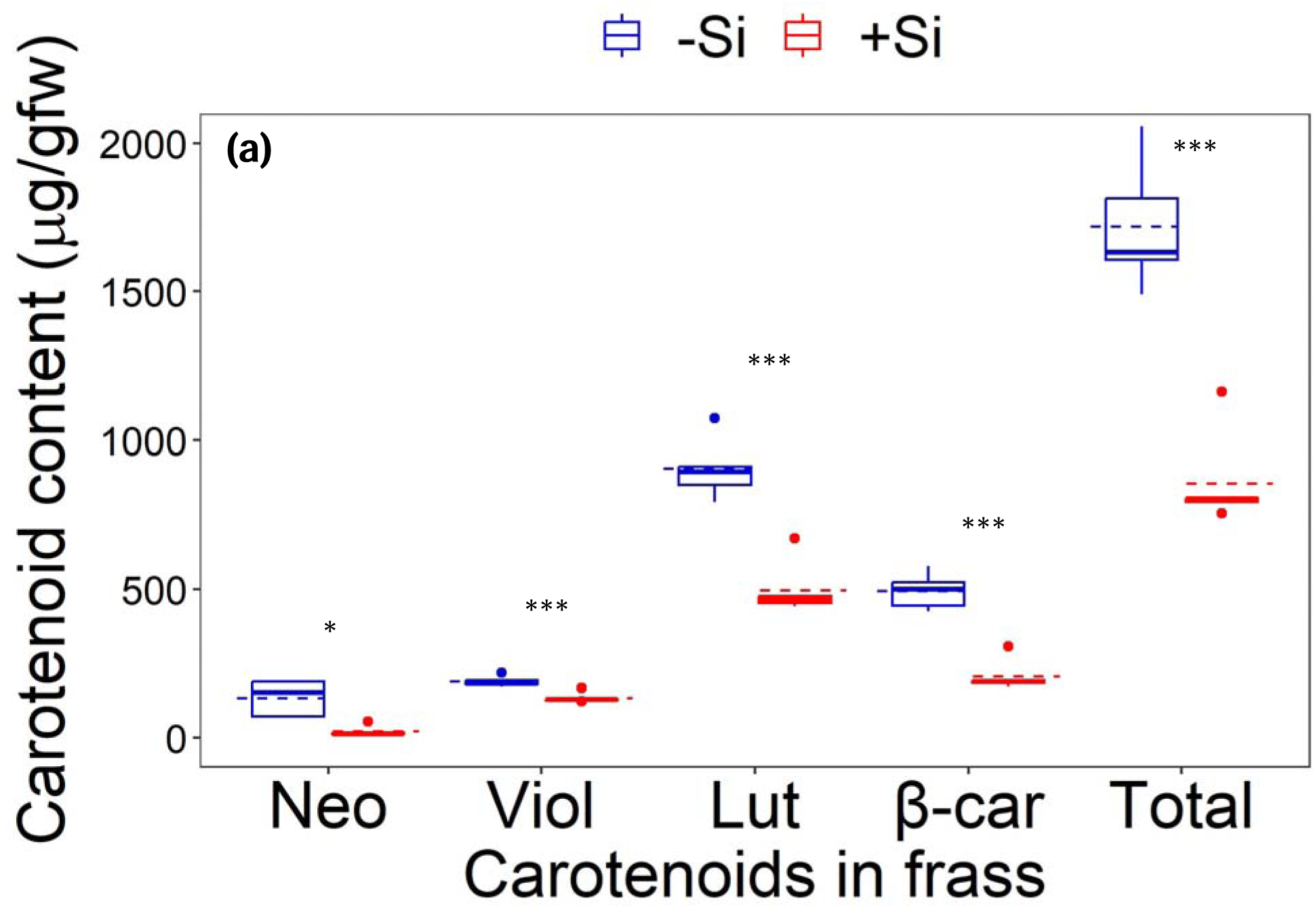
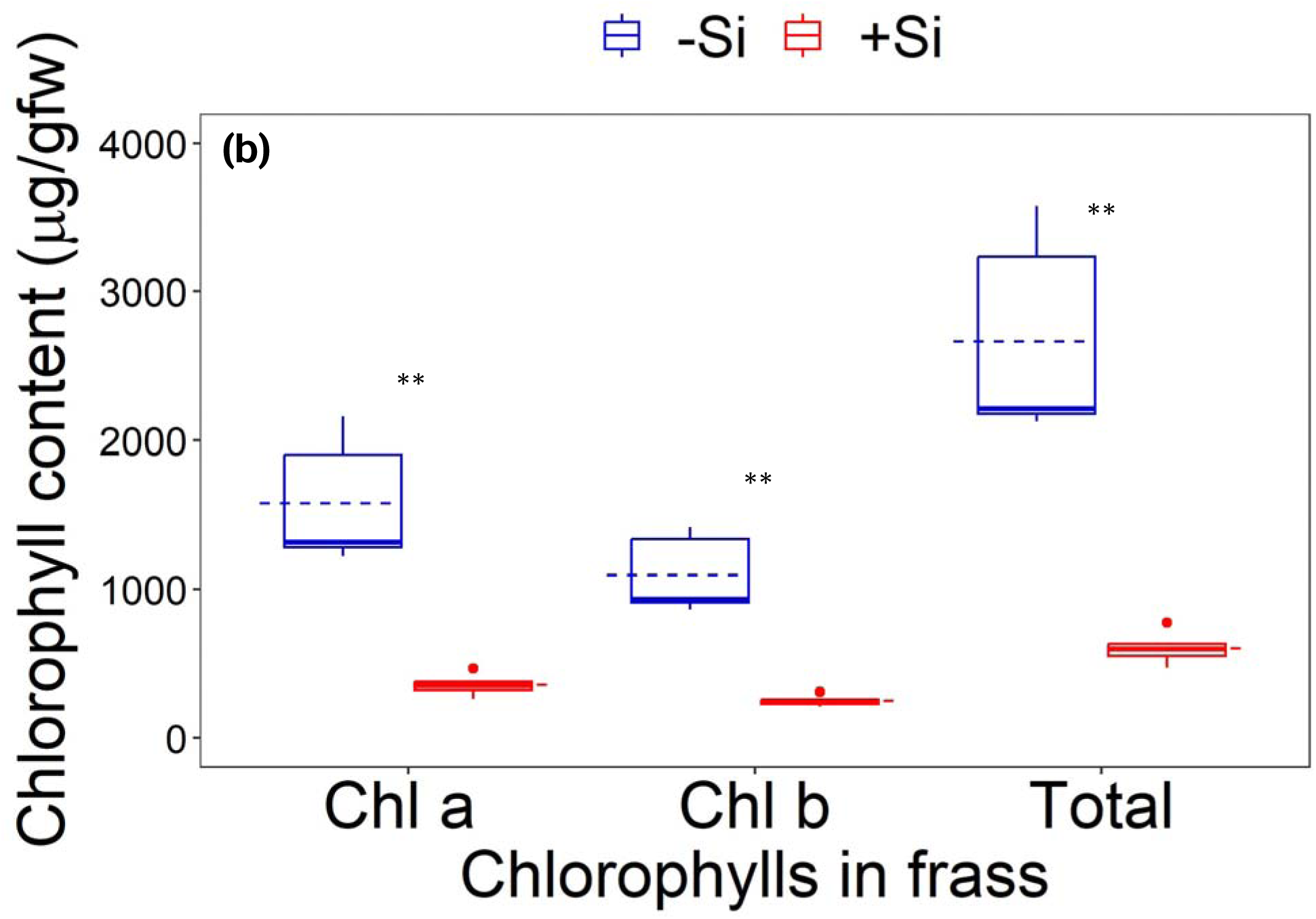
Carotenoid (a) and chlorophyll (b) contents in the frass (µg/g fresh weight) of larvae fed on –Si (silicon-free) and +Si (silicon-supplemented) plants. Boxplots show the interquartile ranges (boxes), non-outlier ranges (whiskers), outliers (solid dots), medians (solid lines) and means (dashed lines). Treatment means were compared using Welch’s *t*-tests. Asterisks indicate levels of statistical significance (* *p* < 0.05, ** *p* < 0.01, *** *p* < 0.001) at a 95% confidence level. Neoxanthin (Neo), violaxanthin (Viol), lutein (Lut), β-carotene (β-car), chlorophyll a (Chl a) and chlorophyll b (Chl b)

### Effects of Si and insect herbivory on carotenoid, chlorophyll and Si levels in the plant

ANOVA models revealed no significant effects of Si, insect herbivory, or their interactions on carotenoid and chlorophyll contents in leaves, except for violaxanthin, where a significant interaction effect was found (Table 2). Under insect-free conditions, pairwise comparisons showed that −Si leaves contained higher levels of total carotenoids (+45%) and chlorophylls (+45%) than +Si leaves (Fig. 4a and 4b; Table 1). For individual carotenoids, violaxanthin (+53%), lutein (+44%) and β-carotene (+49%) were notably higher in –Si leaves than in +Si leaves (Fig. 4a). Similarly, both chlorophyll a (+44%) and chlorophyll b (+48%) were discernibly higher in –Si leaves (Fig. 4b). Following insect herbivory, carotenoid and chlorophyll contents tended to decline in –Si leaves and increase in +Si leaves relative to insect-free conditions, resulting in comparable pigment levels between Si treatments. In response to insect feeding, leaf Si concentrations (% dry weight) significantly increased (+87%) in +Si plants, suggesting the induction of Si defences. Mean Si concentrations were 2.16 ± 0.11% and 3.56 ± 0.27% for insect-free and insect-fed plants, respectively (mean ± SE; Table 1).

**Fig. 4.**
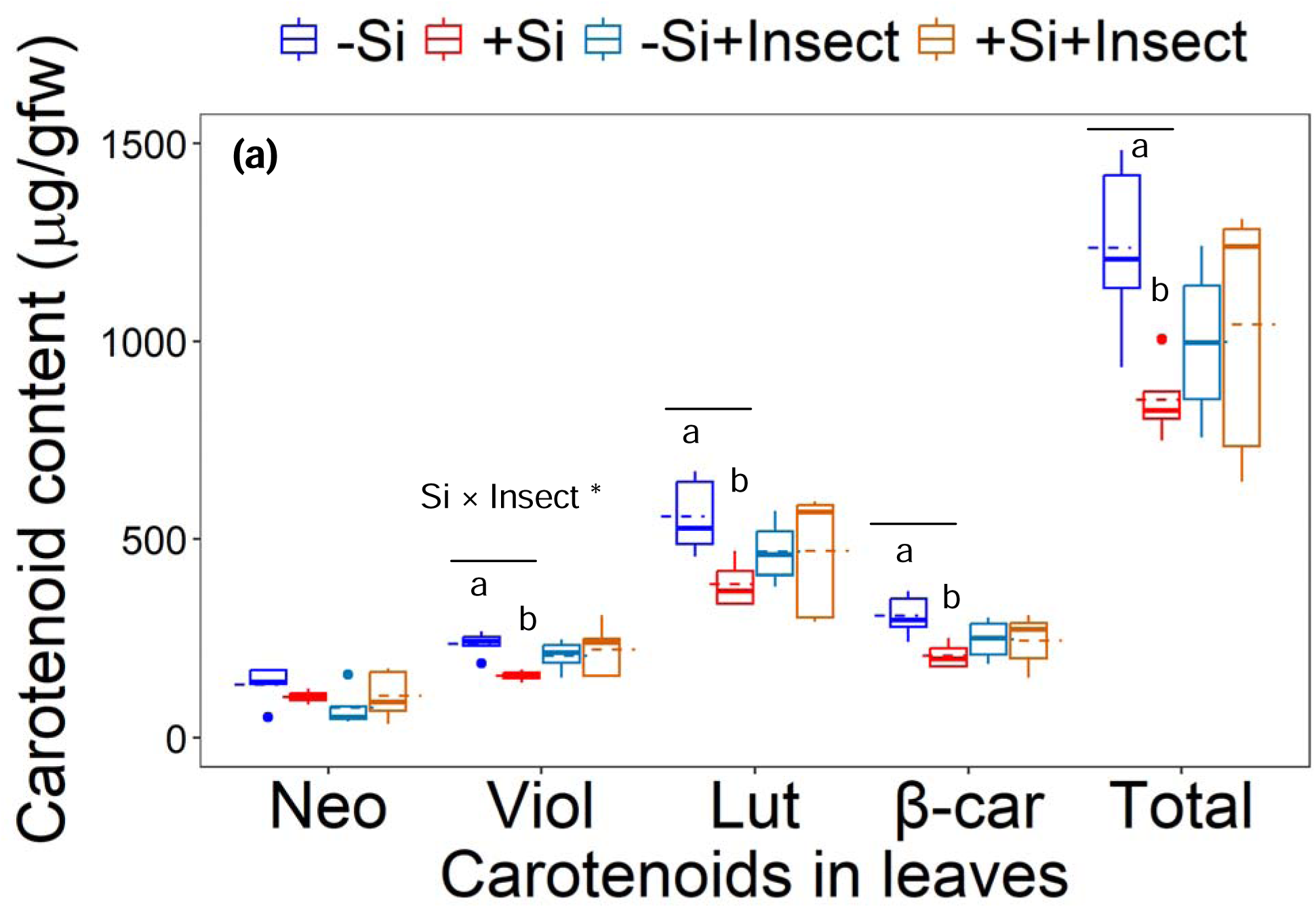
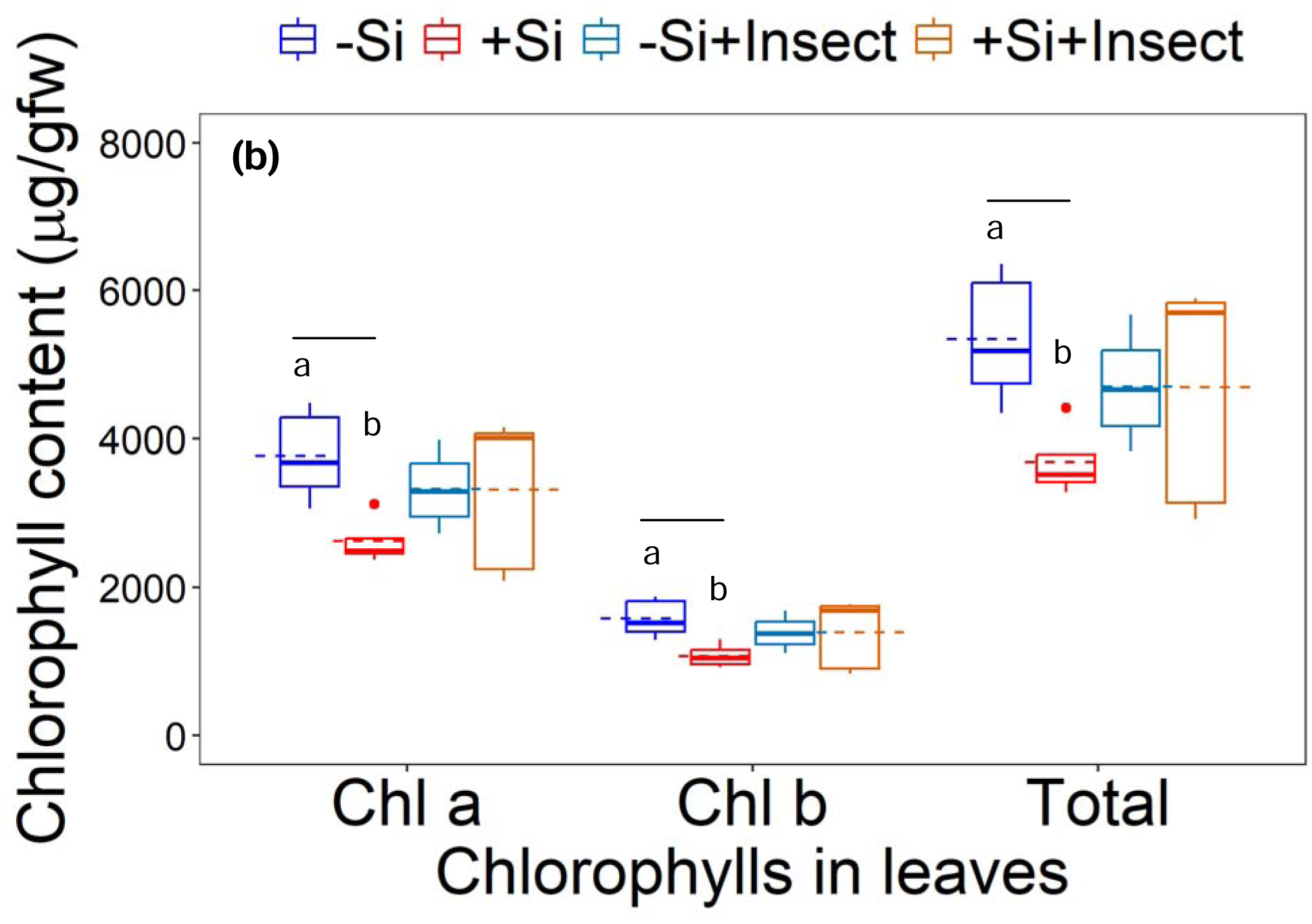
Carotenoid (a) and chlorophyll (b) contents in leaves (µg/g fresh weight) from –Si (silicon-free) and +Si (silicon-supplemented) plants under insect-attacked (+insect) and insect-free conditions. Boxplots show the interquartile ranges (boxes), non-outlier ranges (whiskers), outliers (solid dots), medians (solid lines) and means (dashed lines). The effects of Si and insect attack and their interactions on pigment contents were analysed using two-way ANOVAs. Pairwise comparisons of pigment contents in –Si and +Si plants, under both insect-attacked and insect-free conditions, were performed using Welch’s *t*-tests. The asterisk indicates the level of statistical significance (* *p* < 0.05) at a 95% confidence level. Different letters indicate significant differences between treatment means (*p* < 0.05). Neoxanthin (Neo), violaxanthin (Viol), lutein (Lut), β-carotene (β-car), chlorophyll a (Chl a) and chlorophyll b (Chl b).

**Table 2.**
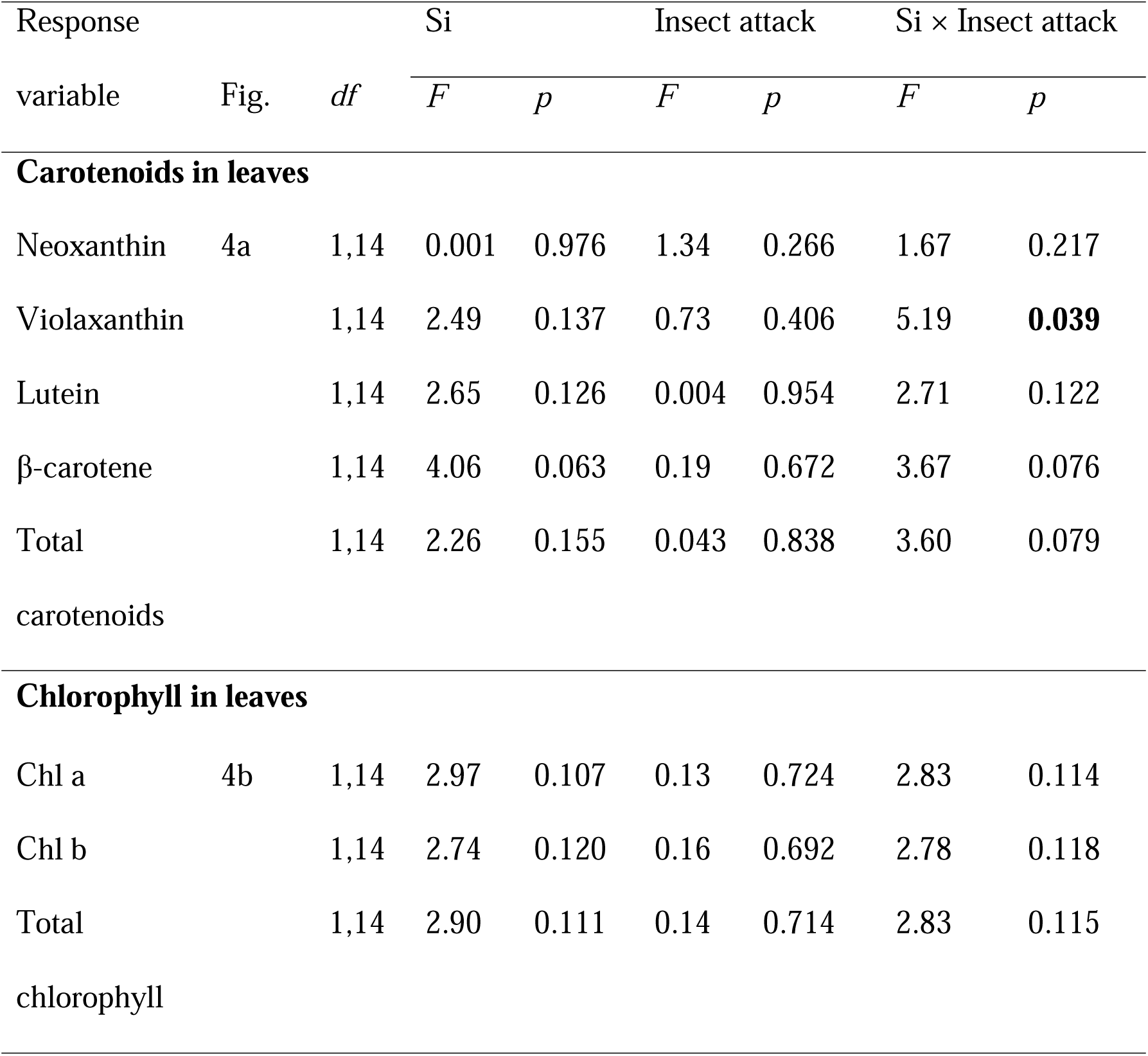
Results of two-way ANOVAs for the effects of Si and insect herbivory on carotenoid and chlorophyll contents in leaves. Statistically significant effect (*p* < 0.05) is indicated in bold.

## DISCUSSION

We provide new evidence that plant Si defences disrupt cryptic colouration in *H. armigera* larvae by reducing carotenoid sequestration into their haemolymph, particularly lutein. This reduction resulted in a conspicuous body-colour polymorphism in larvae: those feeding on −Si plants exhibited cryptic green colouration that blended with leaves, whereas larvae on +Si plants developed contrasting brown colouration. Constitutive levels of all leaf pigments, except neoxanthin, were lower in +Si than in −Si plants. However, following insect herbivory, pigment levels in −Si and +Si plants became comparable.

Previous studies have consistently shown that Si suppresses the performance of chewing insect pests, including *H. armigera* (Biru et al., 2021; Islam et al., 2020; Johnson et al., 2024). Our results add a new dimension by demonstrating that Si not only suppresses larval performance but also disrupts their cryptic colouration by restricting carotenoid acquisition from host plants.

While RGB colour measurements do not directly predict predators’ colour perception, they provide valuable insights into potential predation risk (Welch et al., 2017). The higher colour contrast of larvae on +Si plants could make them more detectible to visually oriented predators, such as spiders, ants and birds (Endler & Mielke, 2005; Greeney et al., 2012). For example, *Trichoplusia ni* larvae with higher colour contrast against their background were five times more likely to be attacked by the predator *Podisus maculiventris* compared to larvae with green cryptic colouration (Welch et al., 2017).

Although −Si and +Si plants synthesised carotenoids (neoxanthin, violaxanthin, β-carotene and lutein) and chlorophylls (a and b) in leaves, larvae only sequestered lutein and β-carotene into their haemolymph, while excreting substantial amounts of pigments regardless of plant Si status.

This suggests that carotenoids are only partially accumulated in insects (Feltwell & Rothschild, 1974; Grayson et al., 1991). Artificial diet-fed larvae lacked pigments entirely, confirming that haemolymph carotenoids were plant-derived. Lutein dominated haemolymph carotenoids, which is consistent with previous studies in Lepidoptera (Cromartie, 1959; Feltwell & Rothschild, 1974; Grayson et al., 1991; Hackman, 1952). In particular, Eichenseer et al. (2002) found that *Helicoverpa zea* larvae preferentially sequestered lutein over other carotenoids, regardless of host plant species. In our study, Si reduced lutein yet increased β-carotene, causing a 44% reduction in total carotenoids in larval haemolymph. Since both β-carotene and lutein can contribute to green colouration via the yellow chromoprotein (Cromartie, 1959; Hackman, 1952), we propose that a minimum threshold of lutein or total carotenoids in the haemolymph is required for larval green colouration.

Leaf pigments, including lutein and β-carotene, were constitutively lower in +Si than in −Si plants under insect-free conditions. Previous studies have reported varied effects of Si on carotenoid and chlorophyll biosynthesis—positive (Ma et al., 2016), neutral (Sienkiewicz-Cholewa et al., 2018) and negative (Harizanova et al., 2019). Although pigment levels equalised between −Si and +Si plants following herbivory, the timing and mechanisms underlying this shift remain unclear. One possible explanation is that under insect-free conditions, Si-supplemented plants invested less in pigment biosynthesis, as silicification can provide metabolic ‘savings’ and reduce dependence on costly chemical defences (Cooke & Leishman, 2011; Epstein, 2009). Under herbivory, +Si plants, benefitting from Si-enhanced structural defences and likely experiencing less damage, may have redirected resources toward pigment biosynthesis to scavenge reactive oxygen species (ROS) (Han et al., 2016; Moreno et al., 2021) and produce volatile apocarotenoids (e.g. β-Ionone) that deter herbivores and attract natural enemies of herbivores (Meng et al., 2023). In contrast, −Si plants, lacking structural defences, likely relied more heavily on pigments for these functions, resulting in pigment depletion.

We consider that the initial lower contents of lutein in +Si leaves contributed to decreased carotenoid sequestration in larvae, with reduced leaf consumption being the primary driver (Islam et al., 2023; Waterman et al., 2021). Our observation aligns with Welch et al. (2017), who showed that *T. ni* larvae with shorter feeding bouts on cabbage leaves accumulated fewer carotenoids and exhibited higher colour contrast. Furthermore, high-silica grasses retained more chlorophyll after digestion by locusts (Hunt et al., 2008), suggesting that Si may hinder pigment digestion, further limiting carotenoid accumulation in larvae. Carotenoids in insects also serve as antioxidants, scavenging ROS produced during immune responses to phytotoxins or pathogens (Heath et al., 2012). Feeding on Si-supplemented plants has recently been shown to enhance phenoloxidase (PO) activity, a key immune enzyme, in *H. armigera* larvae (Islam et al., 2023). Elevated PO activity could deplete carotenoid reserves in larvae, as these pigments are used to mitigate oxidative stress (Cornet et al., 2007).

In conclusion, our study provides novel insights into the ecological significance of Si in plant-insect interactions. *Helicoverpa armigera* larvae feeding on Si-supplemented plants experienced stunted growth and were less adept at accumulating carotenoids, which disrupted their ability to blend with the green leaves. While we did not directly measure predation rates, this disruption in crypsis may increase larval predation risk and influence predator-prey dynamics. Furthermore, Si deposition and Si-induced changes in plant pigment levels could influence larval feeding behaviours and host plant selection, as larvae may seek alternate hosts with greater carotenoid availability or digestibility. Although our findings are based on hydroponically grown plants and *H. armigera*, similar mechanisms may operate in soil-based and natural ecosystems and extend to other pest species that exhibit crypsis. Further studies in more ecologically realistic settings are needed to explore the cascading effects of plant Si defences on insect foraging, cryptic colouration and predator-prey interactions. We suggest that in agricultural ecosystems Si supplementation may provide dual benefits: directly suppressing pest performance and indirectly enhancing biological control by increasing pest visibility to natural enemies.

## AUTHOR CONTRIBUTIONS

**Tarikul Islam:** Conceptualization; Investigation; Formal Analysis; Visualization; Writing – original draft; Writing – review and editing; **Sidra Anwar**: Investigation; Formal Analysis; Writing – review and editing. **Christopher Cazzonelli**: Supervision; Writing – review and editing. **Ben Moore:** Supervision; Writing – review and editing. **Scott Johnson**: Conceptualization; Funding Acquisition; Supervision; Writing – review and editing.

## ACKNOWLEDGEMENTS

We would like to thank Dr Dominika Wojtalewicz for her great help with the HPLC analysis.

## REFERENCES

Ahmad, B., Mutahira, H., Li, M., & Muhammad, M.S. (2019) Measuring focus quality in color space. In 2019 2nd International Conference on Communication, Computing and Digital systems (C-CODE), pp. 115–119.

Alagoz, Y., Dhami, N., Mitchell, C., & Cazzonelli, C.I. (2020). *cis/trans* carotenoid extraction, purification, detection, quantification, and profiling in plant tissues. In Plant and Food Carotenoids (ed. by M. Rodríguez-Concepción & R. Welsch), pp. 145–163. Humana, New York, NY.

Anwar, S., Nayak, J.J., Alagoz, Y., Wojtalewicz, D., & Cazzonelli, C.I. (2022). Purification and use of carotenoid standards to quantify cis-trans geometrical carotenoid isomers in plant tissues. In Carotenoids: Carotenoid and Apocarotenoid Analysis (ed. by E.T. Wurtzel), Vol. 670, pp. 57-85. Academic Press, London.

Baranski, R. & Cazzonelli, C.I. (2016). Carotenoid biosynthesis and regulation in plants. In Carotenoids: Nutrition, Analysis and Technology (ed. by A. Kaczor & M. Baranska), pp. 161–189. Wiley-Blackwell, Oxford, UK.

Bi, J.L. & Felton, G.W. (1995) Foliar oxidative stress and insect herbivory - primary compounds, secondary metabolites, and reactive oxygen species as components of induced resistance. Journal of Chemical Ecology, 21, 1511–1530.

Biru, F.N., Islam, T., Cibils-Stewart, X., Cazzonelli, C.I., Elbaum, R., & Johnson, S.N. (2021) Anti-herbivore silicon defences in a model grass are greatest under Miocene levels of atmospheric CO_2_. Global Change Biology, 27, 2959–2969.

Brkljacic, J., Grotewold, E., Scholl, R., Mockler, T., Garvin, D.F., Vain, P., Brutnell, T., Sibout, R., Bevan, M., Budak, H., Caicedo, A.L., Gao, C.X., Gu, Y., Hazen, S.P., Holt, B.F., Hong, S.Y., Jordan, M., Manzaneda, A.J., Mitchell-Olds, T., Mochida, K., Mur, L.A.J., Park, C.M., Sedbrook, J., Watt, M., Zheng, S.J., & Vogel, J.P. (2011) *Brachypodium* as a model for the grasses: today and the future. Plant Physiology, 157, 3–13.

Cazzonelli, C.I. (2011) Carotenoids in nature: insights from plants and beyond. Functional Plant Biology, 38, 833–847.

Cooke, J. & Leishman, M.R. (2011) Is plant ecology more siliceous than we realise? Trends in Plant Science, 16, 61–68.

Cornet, S., Biard, C., & Moret, Y. (2007) Is there a role for antioxidant carotenoids in limiting self-harming immune response in invertebrates? Biology Letters, 3, 284–288.

Cromartie, R.I.T. (1959) Insect pigments. Annual Review of Entomology, 4, 59-76. Cunningham, J.P. & Zalucki, M.P. (2014) Understanding heliothine (Lepidoptera: Heliothinae) pests: what is a host plant? Journal of Economic Entomology, 107, 881–896.

Czeczuga, B. (1986) The presence of carotenoids in various species of Lepidoptera. Biochemical Systematics and Ecology, 14, 345–351.

Dhami, N., Tissue, D.T., & Cazzonelli, C.I. (2018) Leaf-age dependent response of carotenoid accumulation to elevated CO_2_ in Arabidopsis. Archives of Biochemistry and Biophysics, 647, 67–75.

Dicke, M. (2009) Behavioural and community ecology of plants that cry for help. Plant, Cell & Environment, 32, 654–665.

Eichenseer, H., Murphy, J.B., & Felton, G.W. (2002) Sequestration of host plant carotenoids in the larval tissues of *Helicoverpa zea*. Journal of Insect Physiology, 48, 311–318.

Endler, J.A. & Mielke, P.W. (2005) Comparing entire colour patterns as birds see them. Biological Journal of the Linnean Society, 86, 405–431.

Epstein, E. (2009) Silicon: its manifold roles in plants. Annals of Applied Biology, 155, 155–160.

Feltwell, J. & Rothschild, M. (1974) Carotenoids in thirty eight species of Lepidoptera. Journal of Zoology, 174, 441–465.

Girin, T., David, L.C., Chardin, C., Sibout, R., Krapp, A., Ferrario-Méry, S., & Daniel-Vedele, F. (2014) *Brachypodium*: a promising hub between model species and cereals. Journal of Experimental Botany, 65, 5683–5696.

Gonzalez-Bellido, P.T., Talley, J., & Buschbeck, E.K. (2022) Evolution of visual system specialization in predatory arthropods. Current Opinion in Insect Science, 52, 100914.

Grayson, J., Edmunds, M., Evans, E.H., & Britton, G. (1991) Carotenoids and colouration of poplar hawkmoth caterpillars (*Laothoe populi*). Biological Journal of the Linnean Society, 42, 457–465.

Greeney, H.F., Dyer, L.A., & Smilanich, A.M. (2012) Feeding by lepidopteran larvae is dangerous: A review of caterpillars’ chemical, physiological, morphological, and behavioral defenses against natural enemies. Invertebrate Survival Journal, 9, 7–34.

Guntzer, F., Keller, C., & Meunier, J.-D. (2011) Benefits of plant silicon for crops: a review. Agronomy for Sustainable Development, 32, 201–213.

Hackman, R.H. (1952) Green pigments of the hemolymph of insects. Archives of Biochemistry and Biophysics, 41, 166–174.

Hall, C.R., Mikhael, M., Hartley, S.E., & Johnson, S.N. (2020) Elevated atmospheric CO_2_ suppresses jasmonate and silicon based defences without affecting herbivores. Functional Ecology, 34, 993–1002.

Han, Y., Li, P., Gong, S., Yang, L., Wen, L., & Hou, M. (2016) Defense responses in rice induced by silicon amendment against Infestation by the leaf folder *Cnaphalocrocis medinalis*. PLoS One, 11, e0153918.

Harizanova, A., Koleva-Valkova, L., Stoeva, A., & Sevov, A. (2019) Effect of silicon on the activity of antioxidant enzymes and the photosynthetic rate of cucumber under mite infestation. Scientific Papers. Series A. Agronomy, 62, 519–528.

Harmon, J.P., Losey, J.E., & Ives, A.R. (1998) The role of vision and color in the close proximity foraging behavior of four coccinellid species. Oecologia, 115, 287–292.

Heath, J.J., Cipollini, D.F., & Stireman III, J.O. (2012) The role of carotenoids and their derivatives in mediating interactions between insects and their environment. Arthropod-Plant Interactions, 7, 1–20.

Hodson, M.J., White, P.J., Mead, A., & Broadley, M.R. (2005) Phylogenetic variation in the silicon composition of plants. Annals of Botany, 96, 1027–1046.

Hunt, J.W., Dean, A.P., Webster, R.E., Johnson, G.N., & Ennos, A.R. (2008) A novel mechanism by which silica defends grasses against herbivory. Annals of Botany, 102, 653–656.

Islam, T., Moore, B.D., & Johnson, S.N. (2020) Novel evidence for systemic induction of silicon defences in cucumber following attack by a global insect herbivore. Ecological Entomology, 45, 1373–1381.

Islam, T., Moore, B.D., & Johnson, S.N. (2022a) Plant silicon defences reduce the performance of a chewing insect herbivore which benefits a contemporaneous sap feeding insect. Ecological Entomology, 47, 951–958.

Islam, T., Moore, B.D., & Johnson, S.N. (2022b) Silicon suppresses a ubiquitous mite herbivore and promotes natural enemy attraction by altering plant volatile blends. Journal of Pest Science, 95, 423–434.

Islam, T., Moore, B.D., & Johnson, S.N. (2023) Silicon fertilisation affects morphological and immune defences of an insect pest and enhances plant compensatory growth. Journal of Pest Science, 96, 41–53.

Johnson, S.N., Rowe, R.C., & Hall, C.R. (2020) Silicon is an inducible and effective herbivore defence against *Helicoverpa punctigera* (Lepidoptera: Noctuidae) in soybean. Bulletin of Entomological Research, 110, 417–422.

Johnson, S.N., Waterman, J.M., Hartley, S.E., Cooke, J., Ryalls, J.M.W., Lagisz, M., & Nakagawa, S. (2024) Plant silicon defences suppress herbivore performance, but mode of feeding is key. Ecology Letters, 27.

Lim, U.T. & Ben-Yakir, D. (2020) Visual sensory systems of predatory and parasitic arthropods. Biocontrol Science and Technology, 30, 728–739.

Lyytinen, A. (2001) Insect coloration as a defence mechanism against visually hunting predatorsPhD thesis, University of Jyväskylä, Jyväskylä, Finland.

Ma, D., Sun, D., Wang, C., Qin, H., Ding, H., Li, Y., & Guo, T. (2016) Silicon application alleviates drought stress in wheat through transcriptional regulation of multiple antioxidant defense pathways. Journal of Plant Growth Regulation, 35, 1–10.

Maoka, T. (2020) Carotenoids as natural functional pigments. Journal of Natural Medicines, 74, 1–16.

Massey, F.P. & Hartley, S.E. (2009) Physical defences wear you down: progressive and irreversible impacts of silica on insect herbivores. Journal of Animal Ecology, 78, 281–291.

Meng, K., Eldar-Liebreich, M., Nawade, B., Yahyaa, M., Shaltiel-Harpaz, L., Coll, M., Sadeh, A., & Ibdah, M. (2023) Analysis of apocarotenoid volatiles from lettuce (Lactuca sativa) induced by insect herbivores and characterization of carotenoid cleavage dioxygenase gene. 3 Biotech, 13, 94.

Moran, N.A. & Jarvik, T. (2010) Lateral transfer of genes from fungi underlies carotenoid production in aphids. Science, 328, 624–627.

Moreno, J.C., Mi, J., Alagoz, Y., & Al-Babili, S. (2021) Plant apocarotenoids: from retrograde signaling to interspecific communication. The Plant Journal, 105, 351–375.

Nguyen, K.O., Al-Rashid, S., Clarke Miller, M., Tom Diggs, J., & Lampert, E.C. (2019) *Trichoplusia ni* (Lepidoptera: Noctuidae) qualitative and quantitative sequestration of host plant carotenoids. Environmental Entomology, 48, 540–545.

Pérez-Gálvez, A., Viera, I., & Roca, M. (2020) Carotenoids and chlorophylls as antioxidants. Antioxidants, 9, 505.

R Core Team. (2019) R: A language and environment for statistical computing. R Foundation for Statistical Computing.

Reidinger, S., Ramsey, M.H., & Hartley, S.E. (2012) Rapid and accurate analyses of silicon and phosphorus in plants using a portable X ray fluorescence spectrometer. New Phytologist, 195, 699–706.

Reynolds, O.L., Keeping, M.G., & Meyer, J.H. (2009) Silicon-augmented resistance of plants to herbivorous insects: a review. Annals of Applied Biology, 155, 171–186.

Rogers, D.J. & Brier, H.B. (2010) Pest-damage relationships for *Helicoverpa armigera* (Hübner) (Lepidoptera: Noctuidae) on vegetative soybean. Crop Protection, 29, 39–46.

Schaefer, H.M. & Rolshausen, G. (2006) Plants on red alert: do insects pay attention? Bioessays, 28, 65–71.

Sienkiewicz-Cholewa, U., Sumisławska, J., Sacała, E., Dziągwa-Becker, M., & Kieloch, R. (2018) Influence of silicon on spring wheat seedlings under salt stress. Acta Physiologiae Plantarum, 40, 54.

Teakle, R.E. & Jensen, J.M. (1985). Heliothis punctigera. In Handbook of insect rearing (ed. by P. Singh & R.F. Moore), Vol. 2, pp. 313–322. Elsevier, Amsterdam.

Valadon, L.R.G., Mummery, R.S., Rothschild, M., & Feltwell, J.S.E. (1974) The relationship between carotenoids of certain Lepidoptera and Coleoptera and of their food plants. Biochemical Society Transactions, 2, 1061–1062.

Vicari, M. & Bazely, D.R. (1993) Do grasses fight back? The case for antiherbivore defences. Trends in Ecology & Evolution, 8, 137–141.

Vosteen, I., Weisser, W.W., & Kunert, G. (2016) Is there any evidence that aphid alarm pheromones work as prey and host finding kairomones for natural enemies? Ecological Entomology, 41, 1–12.

Waterman, J.M., Cibils Stewart, X., Cazzonelli, C.I., Hartley, S.E., & Johnson, S.N. (2021) Short term exposure to silicon rapidly enhances plant resistance to herbivory. Ecology, 102, e03438.

Waterman, J.M., Hall, C.R., Mikhael, M., Cazzonelli, C.I., Hartley, S.E., Johnson, S.N., & Cooke, J. (2020) Short term resistance that persists: Rapidly induced silicon anti herbivore defence affects carbon based plant defences. Functional Ecology, 35, 82–92.

Welch, B.J., Obadi, O.M., & Lampert, E.C. (2017) Effects of carotenoid sequestration on a caterpillar’s cryptic coloration and susceptibility to predation. Entomologia Experimentalis et Applicata, 163, 177–183.

Yamasaki, A., Shimizu, K., & Eujisaki, K. (2009) Effect of host plant part on larval body-color polymorphism in *Helicoverpa armigera* (Lepidoptera: Noctuidae). Annals of the Entomological Society of America, 102, 76–84.

Zalucki, M.P., Daglish, G., Firempong, S., & Twine, P. (1986) The biology and ecology of *Heliothis armigera* (Hübner) and *Heliothis Punctigera* Wallengren (Lepidoptera, Noctuidae) in Australia - what do we know? Australian Journal of Zoology, 34, 779–814.

